# Complete genome sequence and metabolic features of *Vreelandella zhaodongensis* BS253: A new isolate from hypersaline lakes from Brazilian Pantanal

**DOI:** 10.64898/2026.01.30.702823

**Authors:** William Lautert-Dutra, Amanda Pasinato Napp, Eduardo Alexandre Back Sivinski, Charley Christian Staats, Francine Melise dos Santos, Clarissa Lovato Melo

**Author notes:** **Corresponding author’s email address** Address correspondence to: Francine Melise dos Santos,; Clarissa Lovato Melo,.

## Abstract

The urgent need for sustainable solutions to mitigate climate change has intensified research into carbon capture, utilization, and storage (CCUS) strategies. Biological approaches, particularly involving extremophilic microorganisms, offer promising alternatives to conventional methods due to their adaptability and potential for bioproduct synthesis. In this study, we report the complete genome sequencing and functional characterization of isolate BS253, derived from a hypersaline alkaline lake in Brazil’s Pantanal region. Using a hybrid sequencing strategy combining Oxford Nanopore long reads and Illumina short reads, we assembled a circular chromosome of 3.76 Mb and identified two plasmids. Phylogenetic and comparative genomic analyses identified the isolate as *Vreelandella zhaodongensis*. Digital DNA-DNA hybridization (dDDH % = 71.4%) and ANI (96,83%) values supported the designation of BS253 as a distinct subspecies of *V. zhaodongensis*. The genome reveals genes associated with salt and alkali tolerance, hydrocarbon and plastic degradation, and the biosynthesis of secondary metabolites. Phenotypically, BS253 is a moderately halophilic, facultatively anaerobic, Gram-negative rod exhibiting biosurfactant activity, with an emulsification index of 51.7% under defined culture conditions. These findings highlight BS253 as a metabolically versatile extremophile with potential applications in different types of industries and biotechnological CCUS systems.

**Importance:** Microorganisms adapted to extreme environments represent an untapped source of biotechnologically valuable traits. *Vreelandella zhaodongensis* BS253, isolated from a hypersaline alkaline lake in the Brazilian Pantanal, expands the known diversity of extremophiles and offers metabolic features with relevance to sustainable bioprocesses. Its complete genome reveals genes involved in salt and alkali tolerance, plastic and hydrocarbon degradation, and the biosynthesis of biosurfactant-like compounds, positioning this strain as a promising chassis for applications in emerging carbon capture, utilization, and storage (CCUS) strategies. The ability of BS253 to produce bioemulsifying molecules under defined nutritional conditions, combined with pathways for degrading recalcitrant pollutants, reinforces its potential for environmentally friendly industrial processes. By characterizing BS253 at the genomic and physiological levels, this work provides foundational information for future exploitation of extremophiles in biotechnological innovations aimed at reducing carbon emissions and supporting circular bioeconomy initiatives.

## Background

The urgency of the climate crisis has heightened the need for effective carbon capture, utilization, and storage (CCUS) strategies to help lower atmospheric CO_2_ levels [1]. A variety of carbon sequestration technologies are currently under investigation globally, which can be divided into two main types: non-biological or biological (microbial or microalgal capture). Non-biological methods frequently face challenges like high operational costs, substantial significant energy requirements, emission of secondary pollutants, scalability issues, and inefficiencies in capture processes [2, 3]. On the other hand, biological technologies present promising alternatives for more sustainable and energy-efficient outcomes.

The isolation of microbial organisms from extreme environments is pivotal for the identification of new strains with potential for ecological and biotechnological applications, including in CCUS strategies [4, 5]. In Brazil, the Pantanal region represents one of the largest tropical wetland ecosystems in the world and hosts a complex mosaic of aquatic environments, including hypersaline lakes. These habitats are influenced by seasonal flooding, high evaporation rates, and fluctuating physicochemical parameters, which contribute to the establishment of highly adapted microbial communities. Such environments harbor unique halophilic and alkaliphilic microorganisms with metabolic versatility. The microbial biodiversity found in hypersaline lakes of the Pantanal therefore constitutes a valuable reservoir for the discovery of novel taxa and bioactive molecules, supporting biotechnological innovation and expanding opportunities for CCUS-related applications [6–9].

In this regard, halophilic environments have shown the presence of distinct taxonomic groups that produce different molecules [10]. For example, organisms within *Halomonadaceae*, the largest family harboring halophilic microorganisms, produce a variety of biotechnologically useful molecules, such as lipids, enzymes, and polysaccharides, that can be employed in several industrial processes [11, 12]. Within this family, the *Vreelandella* genus, recently proposed as one of five new genera from *Halomonas*, currently comprises 39 species. The species show Gram-staining-negative rod cells, aerobic or facultatively anaerobic metabolism, slight to moderately halophilic, alkaliphilic or alkali tolerant, growing at pH values in the range of 5.0–12.0[13].

Whole-genome sequencing methods and computational tools have revolutionized the ways new molecules are characterized. These advancements have allowed the sequencing of diverse environmental organisms, unveiling an extraordinary biological reservoir of biosynthetic potential for secondary metabolites [14]. Secondary metabolites are a broad category of natural products synthesized by organisms, allowing them to adapt to their surrounding environment [15]. For example, certain microorganisms generate biosurfactants, which lower the surface and interfacial tension in biphasic systems, facilitating processes like emulsification, dispersion, and stabilization at interfaces [16, 17]. Moreover, the investigation of secondary metabolites produced by microorganisms offers an environmentally friendly alternative to production chains oriented towards the bioeconomy, contributing to sustainable development on several fronts [18, 19].

Therefore, we characterized the genome of *Vreelandella* strain BS253 in this study. Using long-read sequence technology, we assembled the complete genome of the BS253 strain and explored its metabolic capabilities, the genetic content and biosynthetic capacity for secondary metabolites, and potential biosurfactant production.

## Methods

### Growth conditions, genomic DNA preparation, and Genome sequencing

Strain BS253 was isolated from sediment in a hypersaline alkaline lake in Brazil’s Pantanal Nhecolândia region (19º30’49’’ S, 56º10’01.8’’ W). Growth conditions were performed as previously described [20]. This work is registered in *Sistema Nacional de Gestão do Patrimônio Genético e do Conhecimento Tradicional Associado* (SisGen) under Project code A5C07E4.

For long-read sequencing, pure colonies were collected and suspended in 1ml of Tris-EDTA buffer solution (Sigma-Aldrich; St. Louis, Missouri, EUA) and sent to the Life Sciences Core Facility (LaCTAD; Campinas, Brazil) for DNA extraction, library preparation, and whole genome sequencing. Briefly, QIAcube^®^ (QIAGEN) equipment was used for DNA extraction. Then, DNA quality control was performed using the Agilent TapeStation system (Agilent Technologies; Santa Clara, CA, United States). The resultant DNA was used for whole genome sequencing using nanopore long read technology (Oxford Nanopore Technologies; Oxford, UK). A Nanopore library was constructed using Native Barcoding Kit 96 V14 (SQK-NBD114.96). Library sequencing was done using FLO-PRO114M: PromethION R10 (M Version).

Short-read sequencing was performed as described previously [9, 21]. Briefly, microorganisms were lysed with lysis buffer (NeoSampleX^®^), and DNA was extracted using magnetic beads. Sequencing was done using Illumina NextSeq 1000 (2×150bp, P1-600 Illumina kit) at Neoprospecta Microbiome Technologies (Florianópolis, Santa Catarina, Brazil).

### Genome assembly and plasmid assembly

The quality and filtering of PE raw reads were analyzed using *FastQC* v0.11.9 and *fastp* v0.23.4 (*trim-poly-g -detect-adapter minlenght 90bp*) [22, 23]. Long-reads were filtered by quality using *Filtlong* v0.2.1 [24] using short reads as *k*-mer matches to the reference to better filter by quality (‘*PE short-reads’* + *--min_length 1000 --keep_percent 90*). Then, filtered nanopore reads were assembled using *Trycycler* v0.5.5, a consensus long-read assembler for bacterial genomes [25], which relies on multiple independent long-read assemblies of the same genome and produces a final consensus *de-novo* long-read genome assembly. We choose *Flye* v2.9.5-b1801 [26], *Minipolish* (*miniasm_and_minipolish*.*sh*) v0.1.2 [27], and *Raven* v1.8.3 [28] assemblies for multi-assembly. Assemblies were manually curated using *Bandage* v0.8.1 [29] and fragmented genomes were discarded. After the sequential steps, the final consensus was polished using *Medaka* v2.0.1 and a short-read polishing using *Polypolish* v0.6.0 and *Pypolca* v0.3.1 [30, 31]. The presence of plasmids was evaluated using *Plassembler* v1.6.2 [32]. This method uses a hybrid approach to remove chromosomal reads and automatically assemble plasmids. Assembled plasmids were annotated following the same approach of the genome sequence (Table 1).

**Table 1.**
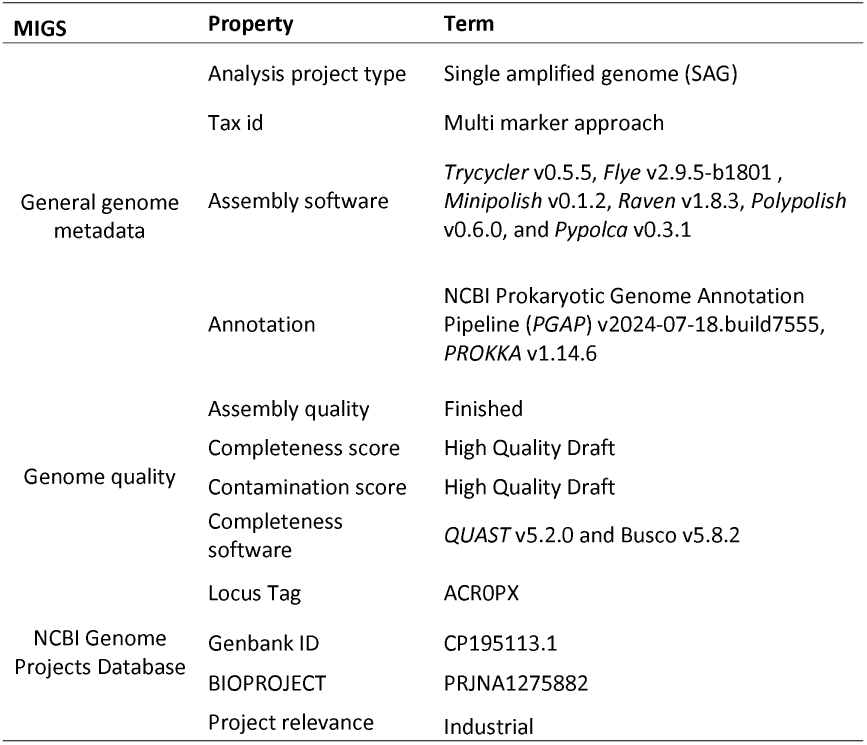
Project information.

### Genome annotation

Different annotation packages were implemented to uncover the genetic characteristics of the BS253 genome. Genome sequencing annotation was done using *PROKKA* v1.14.6 and *PGAP* v*2024-07-18*.*build7555* (NCBI Prokaryotic Genome Annotation Pipeline) pipelines [33, 34]. To assess the genome’s completeness, we used the Benchmarking Universal Single-Copy Orthologue (*BUSCO*) tool v5.8.2 to provide completeness and redundancy analysis related to expected gene content in the *de novo* assembled genome and annotated gene set [35]. Assembly quality and coverage were evaluated using *QUAST* v5.2.0 [36]. Finally, we investigated the antibiotic resistance genes presented in the BS253 genome using the Comprehensive Antibiotic Resistance Database (CARD) [37].

### Phylogenetic and genome similarity analysis

Phylogenomic analysis was then used to assess the evolutionary relationship. First, after-tax identification by *PGAP* (-*-taxcheck*) in the annotation step, we downloaded RefSeq reference genomes from the assigned genes (*Vreelandella*) (Additional file: Table S1) using the *datasets* v16.40.1 tool [38]. Then, the BS253 and reference genomes were mapped to the *Phylophlan* database [39] using *DIAMOND* v2.1.11.165 [40]. Multi-fasta mapped files for core genes were aligned and trimmed using *MUSCLE* (v5.3.linux64) and *trimAL* (v1.5.rev0 build[2024-05-27]) [41, 42]. These files were used to compute independent gene trees using *RaxML* v8.2.12 (ML + LG matrix-based model) for all genes [43]. The final trees were trimmed using *TreeShrink* v1.3.9 with different false positive error rates (*-q* “0.05, 0.10, 0.10”) options. After, each final evolutionary species tree was generated using *ASTRAL* v5.7.8 [44]. The ASTRAL normalized score was used to choose the best species tree. The *Carnimonas nigrificans* (Accession: GCF_000526695.1) genome was used as an outgroup [45].

To determine if the BS253 was related to the previously described species and following the minimal standards for prokaryote taxonomy [46], Average Nucleotide Identity (ANI) was calculated for the BS253 and reference sequences (Table S1) with *fastANI* v1.34 [47]. Additionally, the BS253 genome sequence was uploaded to the TYGS web server for dDDH (digital DNA-DNA hybridization) computed with GGDC (Genome-to-Genome Distance Calculator) [48, 49].

### Microbial secondary metabolites and special metabolism prediction

The presence of specialized metabolism linked to hydrocarbon degradation, plastic degradation, and biosurfactant production was done using *HADEG* (*HADEG_protein_database_231119*.*faa*) [50, 51]. Also, we characterized the presence of Toxin-antitoxin (TA) loci presented in the genome using the TADB database web server [52].

The presence of biosurfactant and potential production was evaluated using experimental methods. First, the microorganism BS253 was initially cultured in saline broth, for 24 h [53]. Subsequently, the cells were harvested by centrifugation at 10.000 rpm for 10 min and washed twice with sterile saline solution (0.9% NaCl). The washed cells were then resuspended in MSM medium [54] supplemented with 2% glucose as a carbon source. The cell suspension [1% (v/v)] was used as the inoculum for biosurfactant induction. The experiment was incubated for 5 days at 28°C and 180 rpm. The ability to produce biosurfactants was evaluated through the emulsification activity, drop collapse and oil spreading assays [55–57]. The negative control consisted of sterile culture medium, and the positive control comprised 0.5% SDS.

### Microbial phenotypic and biochemical features

The physiological and biochemical properties of strain BS253 were characterized following standard microbiological protocol [58]. Gram staining and cellular morphology were assessed by light microscopy (Nikon Eclipse Si) using cells cultivated in a modified LB medium (1.0% w/v tryptone and 0.5% w/v yeast extract, supplemented with 1%, 5% or 10% NaCl). Colony morphology was also examined under these conditions. Biochemical profiling was conducted through a combination of conventional phenotypic assays, following established methodologies [59] and the API miniaturized system protocol (bioMérieux). Growth optimization was evaluated in a modified LB medium supplemented with different NaCl concentrations (0, 3, 4, 5, 10, 12.5, 15 and 20% w/v). To determine the pH range compatible with growth, the culture medium was adjusted to pH values from 3 to 13 using specific buffering systems [60]. The thermal tolerance range was evaluated by incubating the cultures at 10, 20, 30, 40, and 50°C.

## Results

### BS253 complete genome properties

The BS253 genome was sequenced by combining Illumina and Nanopore sequencing technologies. Using multiple independent long-read assemblies of the same genome, we produced a final consensus *de-novo* long-read genome assembly with 3,765,793 bp (Table 2). This complete genome was distributed in a long circular chromosome (Figure 1) and with G+C content, N90, and N’s bases equal 53.35%, 1, and 0, respectively (Table 2). The trimmed reads represent approximately 529-fold and 237-fold coverage of the genome from Illumina and Nanopore reads, respectively (Table 2). The BS253 genome also showed one copy each of two plasmids, one with 28,749 bp and the other with 6,228 bp (Table 2). The coverage of both plasmids aligned with the chromosome coverage for Illumina and Nanopore reads (Table 2).

**Table 2.**
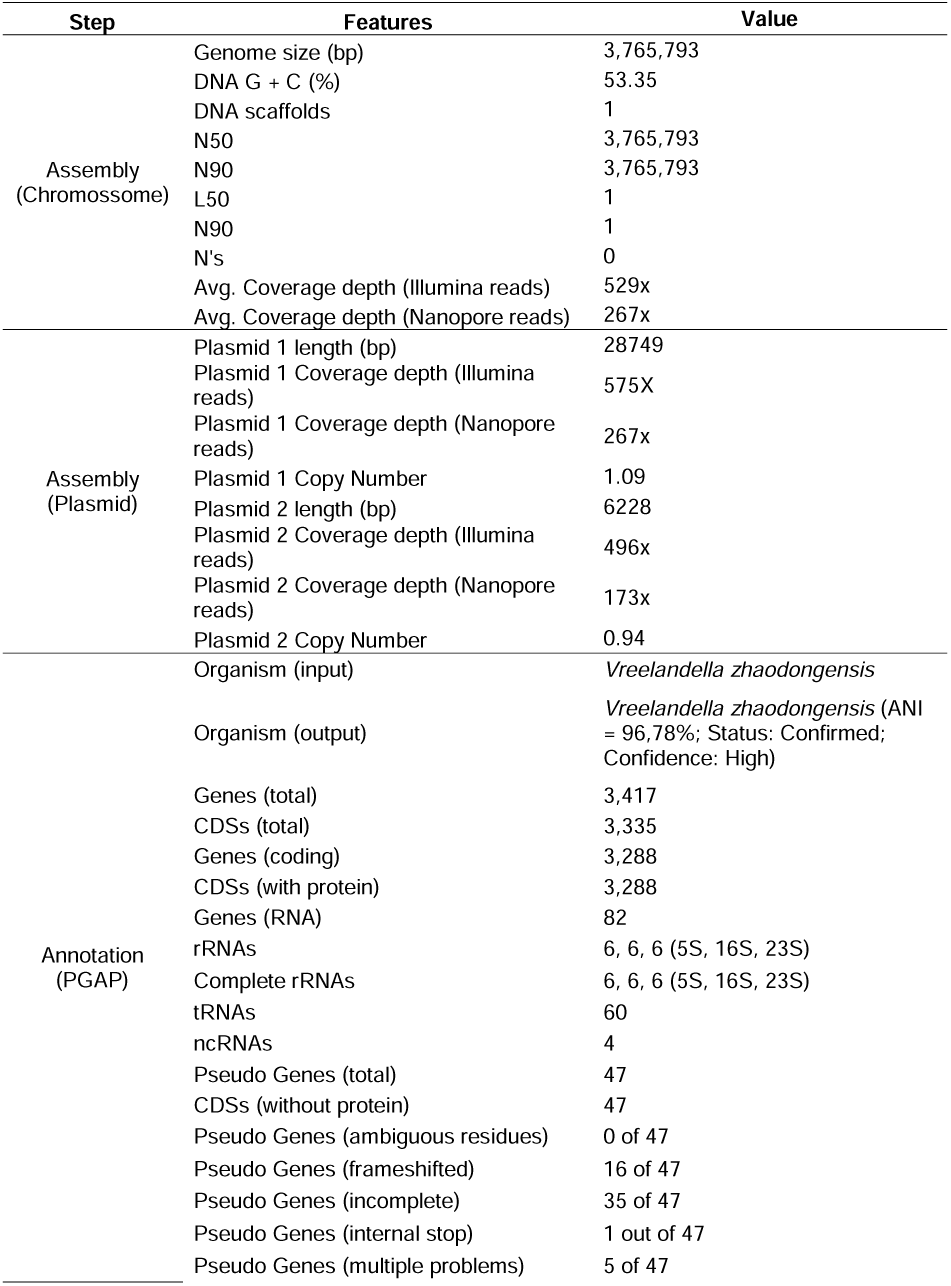
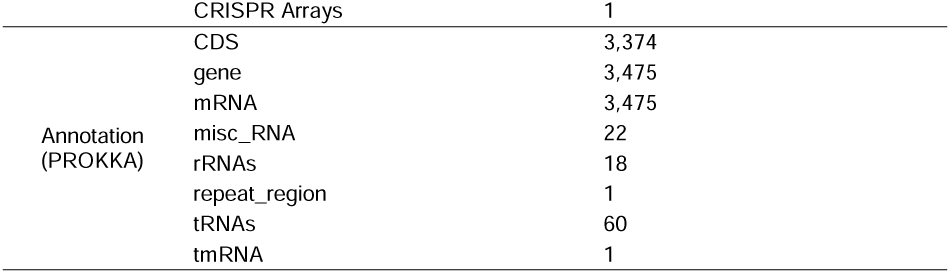
Genome statistics. The summary highlights the major statistics for each step from *de novo* genome assembly, plasmid assembly to genome annotation.

**Figure 1.**
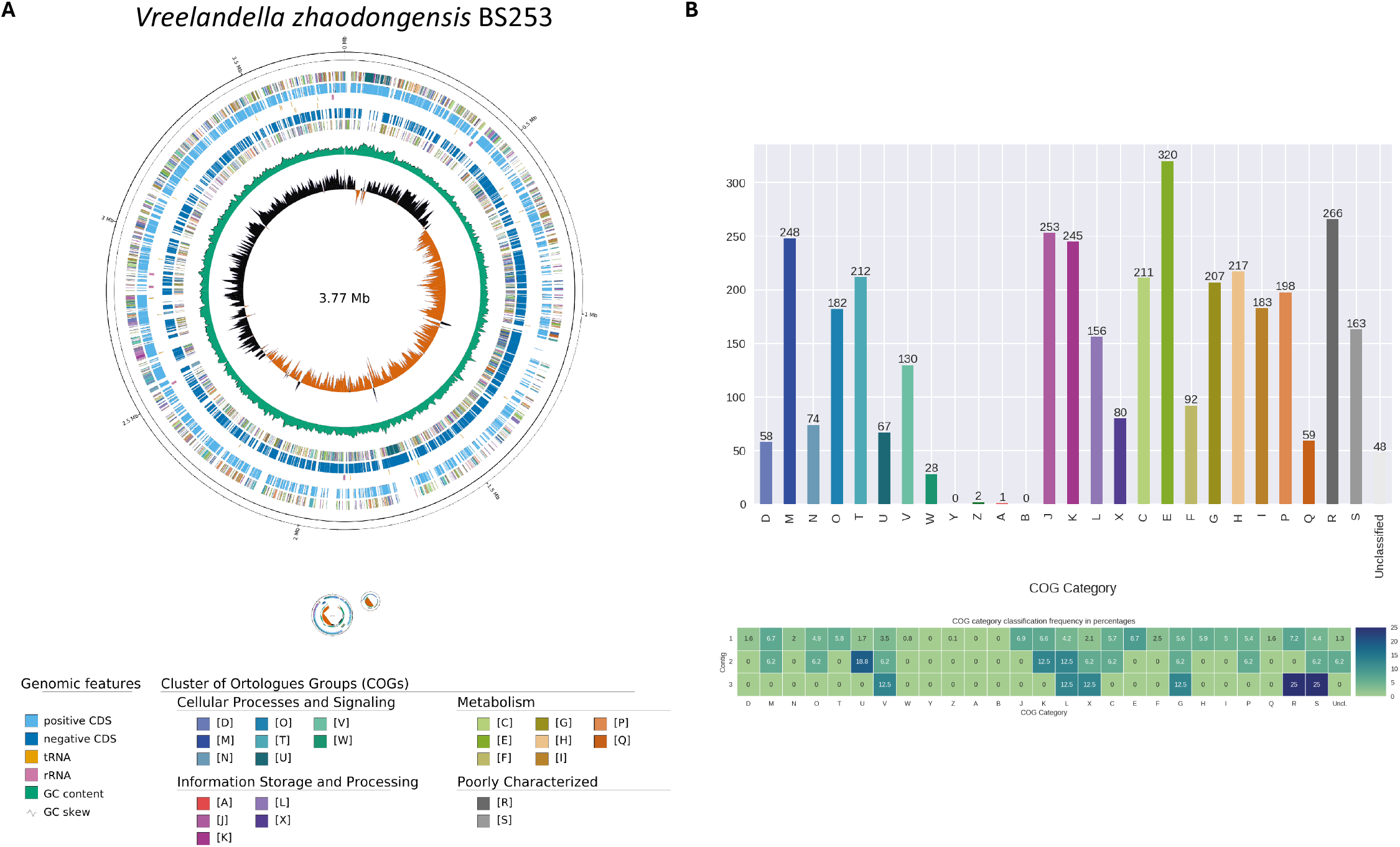
Figure 1. Circular representation of the complete de novo assembled genome of Vreelandella zhaodongensis BS253. **(A)** Overview of the genome of isolate BS253 showing COG functional categories displayed for the metabolism coding region predicted in the assembled genome. **(B)** A histogram displaying COG classification categories of every CDS named. CDS, Coding sequence; tRNA, transfer RNA; rRNA, ribosomal RNA; GC, guanine-cytosine; D, Cell cycle control, division, chromosome partitioning; M, Cell wall/membrane/envelope biogenesis; N, Cell motility; O, Post-translational modification, protein turnover, chaperones; T, Signal transduction mechanism; U, Intracellular trafficking, secretion, and vesicular transport; V, Defense mechanism; W, Extracellular structures; Y, Nuclear structure; Z, Cytoskeleton; A, RNA processing and modification; B, Chromatin structure and dynamics; J, Translation, ribosomal structure, and biogenesis; K, Transcription; L, Replication, recombination, and repair; X, Mobilome: prophages, transposons; C, Energy production and conversion; E, Amino acid transport and metabolism; F, Nucleotide transport and metabolism; G, Carbohydrate transport and metabolism; H, Coenzyme transport and metabolism; I, Lipid transport and metabolism; P, Inorganic ion transport and metabolism; Q, Secondary metabolites biosynthesis, transport, and metabolism; R, General function prediction only; S, Function unknown; COG, Clusters of Orthologous Groups. Generated by GenoVi (https://github.com/robotoD/GenoVi).

The two different annotation software used produced similar and complementary results. In the *PGAP* annotation, the BS253 genome encodes 3,335 CDS, 3,288 genes, and 60 tRNAs—the rRNAs segments comprised six copies of each 5S, 16S, and 23S. While using the *PROKKA* pipeline, the genome of BS253 encodes 3,374 CDS, 3,475 mRNAs, 18 rRNAs, and 60 tRNAs. The *PGAP* annotation also added GO annotation for annotated genes. The genome showed 4830 GO related to molecular function (MF, 2632; Additional file: Figure S1a), biological process (BP, 1606; Additional file: Figure S1ab), and cellular component (CC, 592; Additional file: Figure S1ac).

The Clusters of Orthologous Groups of proteins (COGs) for each CDS showed high frequency for amino acid transport and metabolism (E), general function (R), translation, ribosomal structure, and biogenesis (J), cell wall/membrane/envelope biogenesis (M), and transcription (K) (Table 3; Figure 1b). The plasmids showed the presence of COGs related to defense mechanism (V), transcription (K), replication, recombination, and repair (L), mobilome, prophages and transposons (X), energy production and conversion (C), carbohydrate transport and metabolism (G), and inorganic ion transport and metabolism (P) (Table 3).

**Table 3.**
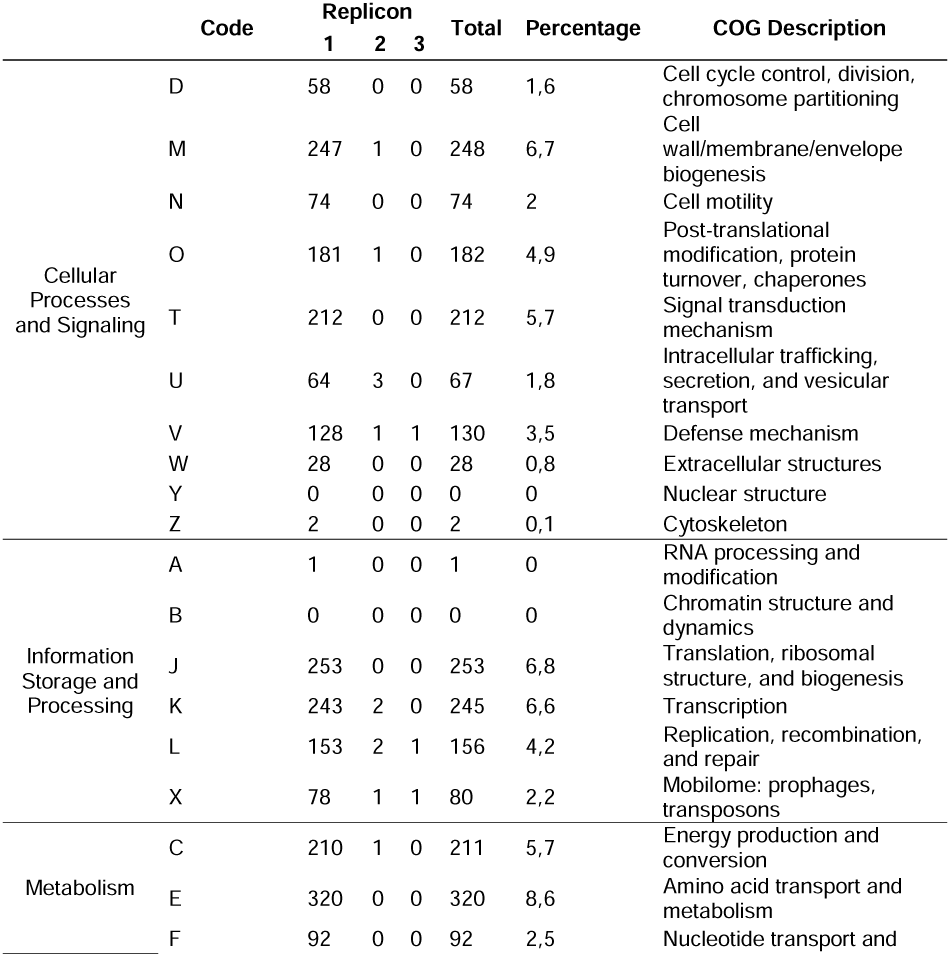
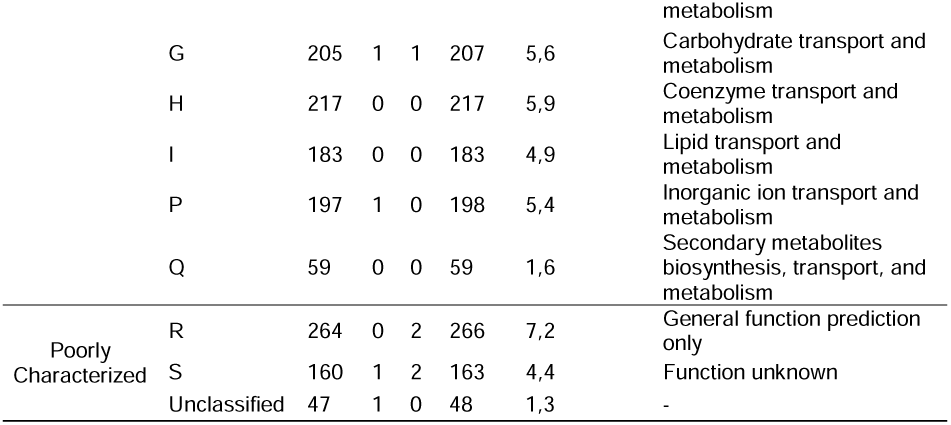
The number of genes associated with general COG functional categories. COG, Clusters of Orthologous Groups. Generated by *GenoVi* (https://github.com/robotoD/GenoVi).

Distinct classes of antibiotic resistance genes are present in the BS253 genome. Small multidrug resistance (SMR), antibiotic efflux pump (*qacG*), vanH gene in vanO cluster (*vanH*), and resistance-nodulation-cell division (RND) antibiotic efflux pump (*rsmA*) were annotated. The resistance mechanisms of these classes are linked to the mechanism of antibiotic efflux and antibiotic target alteration and drugs like disinfecting agents and antiseptics, glycopeptide antibiotic, fluoroquinolone antibiotic, diaminopyrimidine antibiotic, and phenicol antibiotic (Table S2). Then, we performed the BUSCO to assess the completeness of the *de novo* assembled genome and annotated gene set using the *halomonas*_*odb12* (Creation date: 2024-11-14, number of genomes: 102, number of BUSCOs: 1250). Among the conserved orthologs, 98.7% were retrieved in both *de novo* assembled genome and annotated gene set (C:98.7% [S:98.2%, D:0.6%], F:0.4%, M:0.9%, n:1250; Figure S2). This result indicates that the *de novo* assembled genome and coding region of the assembly are highly complete.

### Phylogenetic analysis and evolutionary history of Vreelandella zhaodongensis BS253

We used a multi-marker approach to determine if the BS253 was related to the previously described species from the genus *Vreelandella*, which was the genera indicated after *PGAP* annotation (-*-taxcheck*). The BS253 *de novo* assembled genome best matched *Vreelandella zhaodongensis* (taxid = 1176240) with high confidence. Further ANI analysis between BS253 and *Vreelandella* reference genomes confirmed this classification, supported by an ANI value of 96,83% between BS253 and *V. zhaodongensis* (GCF_013415115.1). The dDDH value confirmed this assumption and showed values above the minimal threshold for a new species (dDDH % = 71.4%). The phylogenetic tree confirmed these results and placed BS253 within *V. zhaodongensis* (Figure 2), with *V. alkaliphila, V. Venusta*, and *V. andesensis* as the nearest related species.

**Figure 2.**
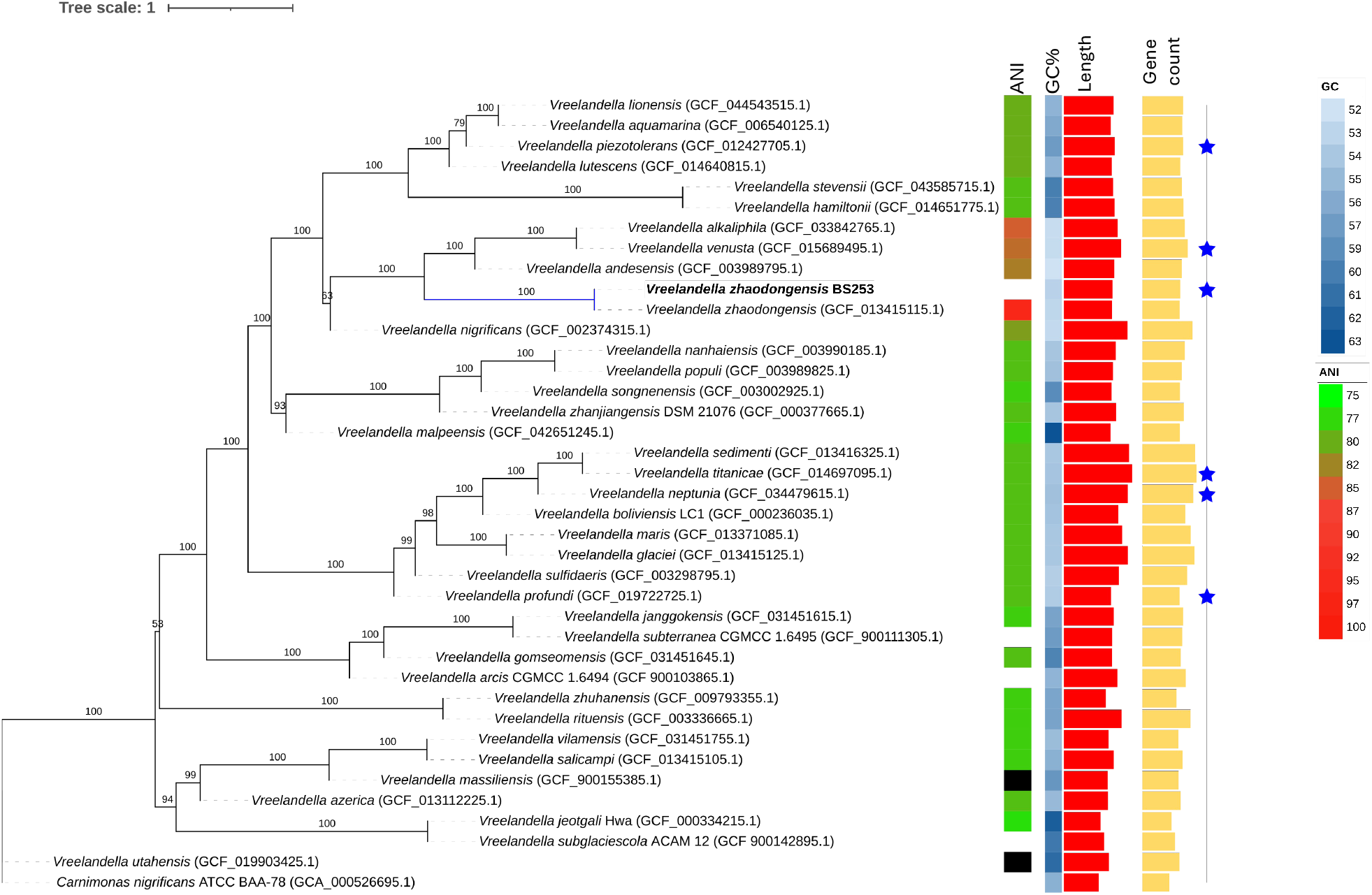
The phylogenetic species tree and correct placement of the isolate BS253 in the *Vreelandella*. Gene trees were inferred using *RaxML* (ML + LG matrix-based model) for 400 genes in *Phylophlan db*. The final trees were trimmed using *TreeShrink* v1.3.9. The evolutionary species tree was generated using *ASTRAL*. The bootstrap value is shown next to the branches. The *Carnimonas nigrificans* genome was used as an outgroup. ANI values of 253 against reference genomes are shown in the heatmap. Genome statistics are displayed, from left to right: Total length (bp), GC% content, and Gene count. Genomes that are classified as complete sequences are marked with blue stars.

### Phenotypic and biochemical characterization

Cells of strain BS253 were identified as Gram-negative, facultative anaerobic, and non-motile rods. Colony morphology varied slightly with salt concentration but was consistently described as cream or beige-colored, circular, convex, exhibiting an entire margin, smooth surface, opaque consistency, and a dull appearance. Colony diameters varied according to NaCl concentration in the modified LB medium: 2.3–3.5 mm at 1% NaCl, 1.8–2.8 mm at 5% NaCl, and 0.6–1.2 mm at 10% NaCl. Cell size was also influenced by salinity, with rod-shaped cells measuring 1.2–2.0 × 0.5–0.8 µm in 1% NaCl, 1.3–2.4 × 0.6–0.9 µm in 5% NaCl, and 1.0–1.6 × 0.5–0.7 µm in 10% NaCl.

The physiological and biochemical characteristics of strain BS253 are detailed in Table 4. The isolate showed oxidase-positive and catalase-negative results. BS253 is classified as moderately halophilic, exhibiting growth in (i) NaCl concentrations ranging from 3% to 12.5% (optimum 3% and 5%), (ii) at temperatures of 10 to 30 °C (optimum between 20°C and 30°C), and (iii) pH range 7 to 11 (optimum pH 8) (Figure 3). Along with this, the strain reduces nitrate, but tested negative for indole production, Voges-Proskauer, and ONPG assays, under aerobic conditions. It hydrolyzed esculin, but showed no hydrolytic activity against gelatin, starch, or Tween 80, and did not produce hydrogen sulfide (H_2_S). The strain also showed no detectable activity for tryptophan deaminase, arginine dihydrolase, phenylalanine deaminase, lysine decarboxylase, ornithine decarboxylase, or urease. Fermentative/oxidative capacity was detected by the production of acids from D-glucose, sucrose, L-arabinose, maltose, L-rhamnose, D-mannitol, sorbitol, and inositol. At the same time, lactose was not utilized as a fermentable substrate. The following compounds were used as sole sources of carbon and energy: N-acetyl-glucosamine, maltose, and citrate.

**Table 4.**
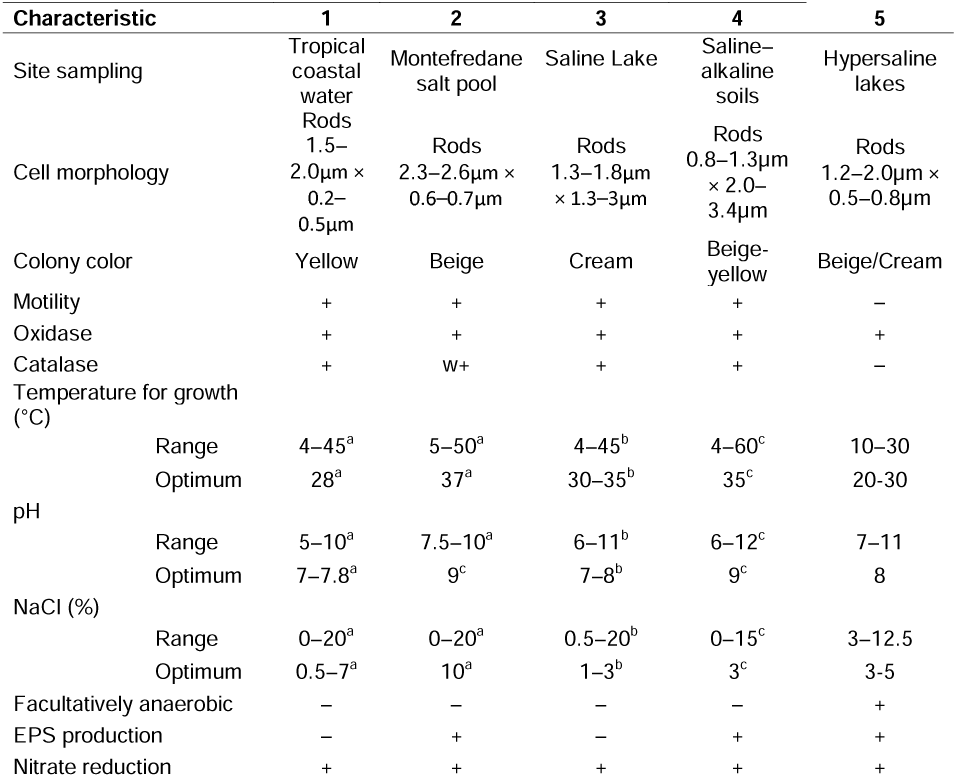
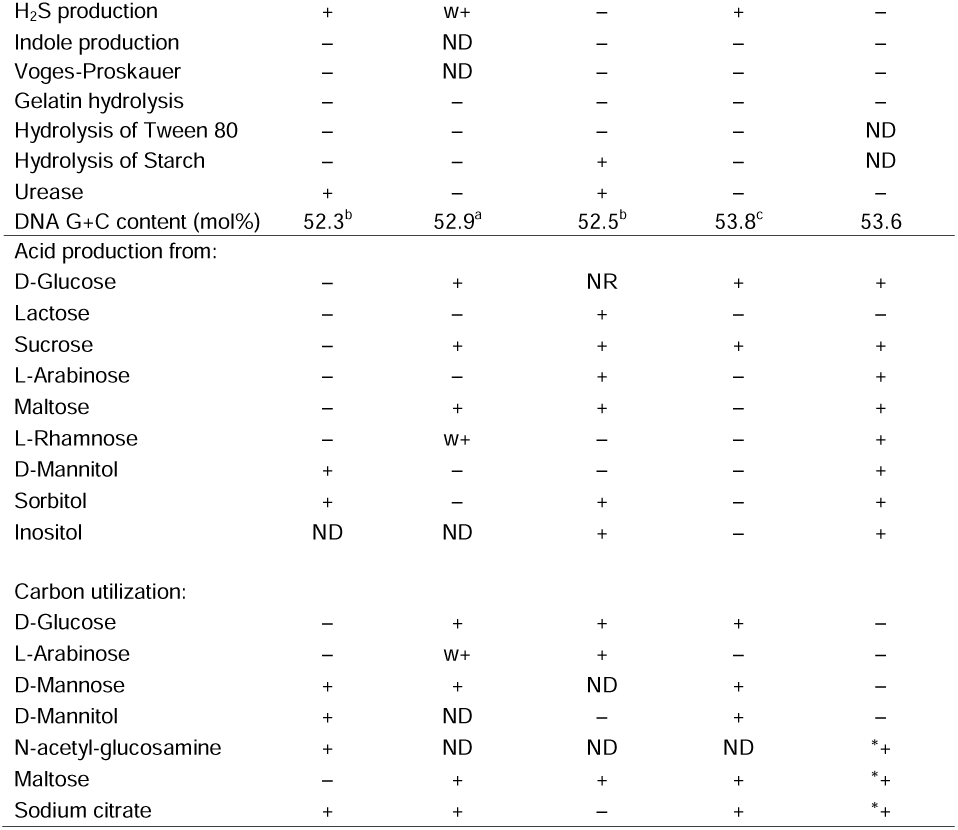
Characteristics that differentiate strain BS253 from closely strains of species of the *Vreelandella*. Strain: 1, *V. venusta* (GCF_015689495.1) [78–81]; 2, *V. alkaliphila* (GCF_033842765.1) [82]; 3, *V. andesensis* (GCF_003989795.1) [83]; 4, *V. zhaodongensis* (GCF_013415115.1) [60]; 5, *V. zhaogonensis* BS253. All data for BS253 strain were generated in this study. +, positive; -, negative; w+, weakly positive, *+ 96h positive, ND, not determined.

**Figure 3.**
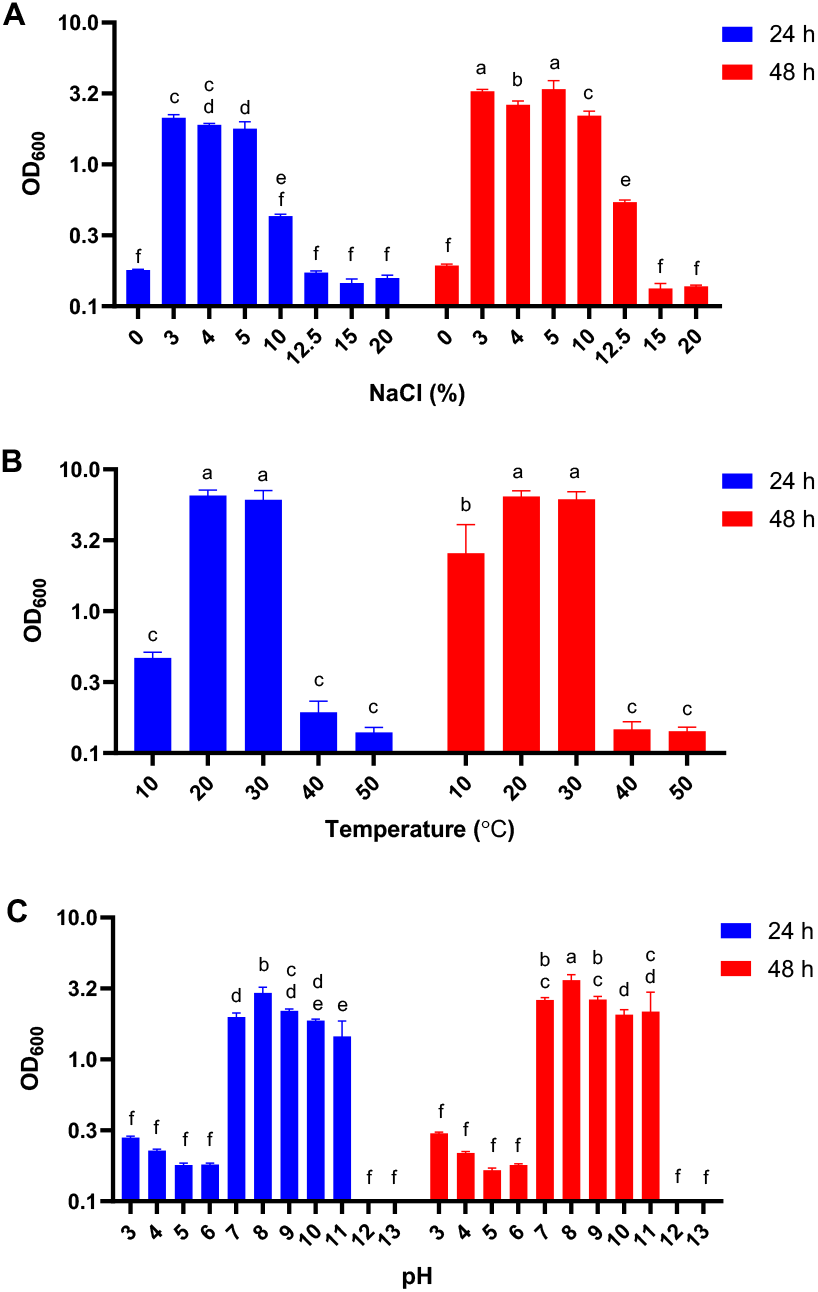
Effect of physicochemical conditions on the growth of strain BS253 after 24 h and 48 h of incubation under different parameters. **(A)** NaCl concentrations (%), **(B)** temperature (°C), and **(C)** initial pH values. Bars represent the mean values, and different letters indicate statistically significant differences (*p* < 0.05) according to Tukey’s test.

### Secondary metabolites and special metabolism

According to HADEG analysis, *V. zhaodongensis* BS253 has routes for key pathway groups related to (i) alkanes, (ii) aromatics and (iii) plastic degradation. The specific subpathways identified include: the Finnerty pathways(*ahpC* and *ahpF*), hydrocarbon uptake (*blc*), catechol degradation (*pcaF* and *pcaJ*), gentisate degradation (*nagL*), homogentisate degradation (*hmgA*), naphthalene degradation (*nahF*), phenylacetate degradation (*paaA, paaB, paaC*, pad, and *paaK*), protocatechuate degradation (*pcaG* and *pcaH*), toluene degradation (*pobA*), PBAT/PET degradation (*est*) (Table 5 and Figure 4).

**Table 5.**
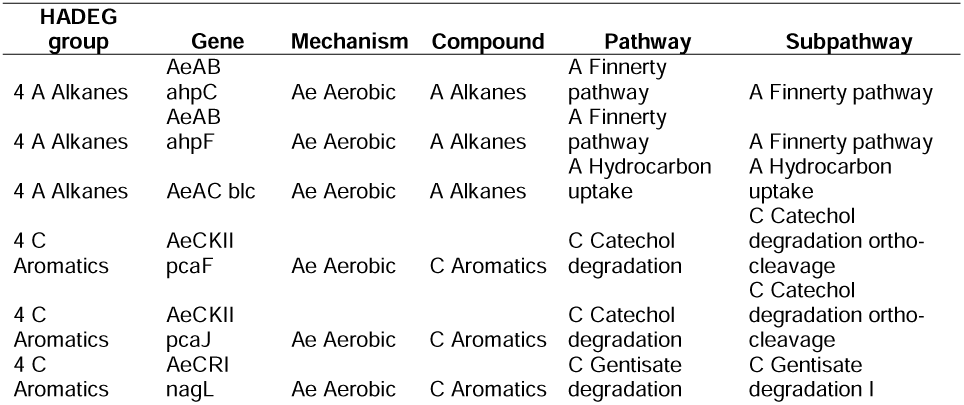
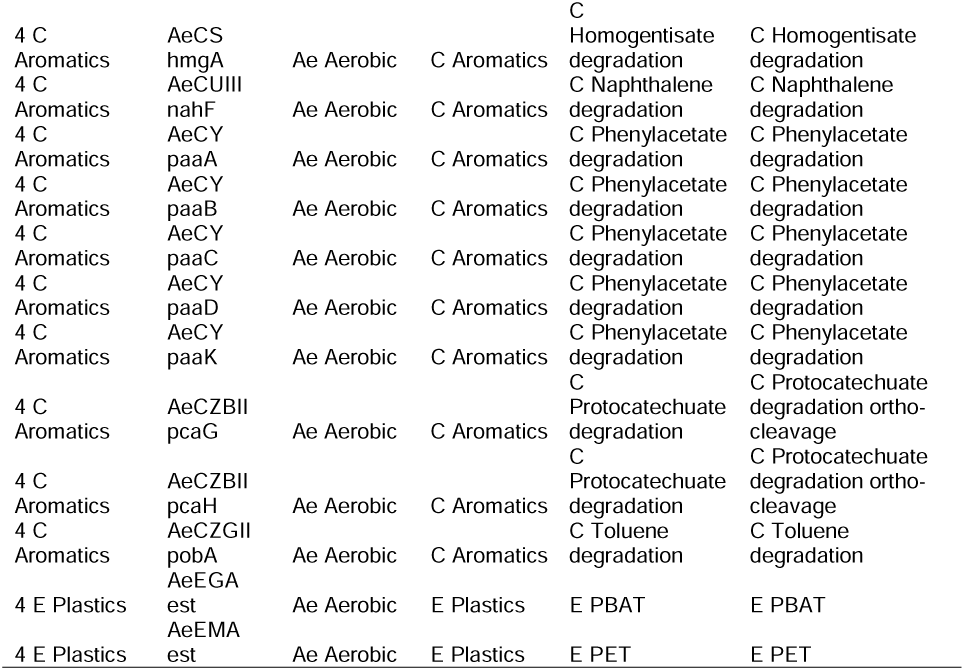
Degradation pathways and subpathways are present in the BS253 genome.

**Figure 4.**
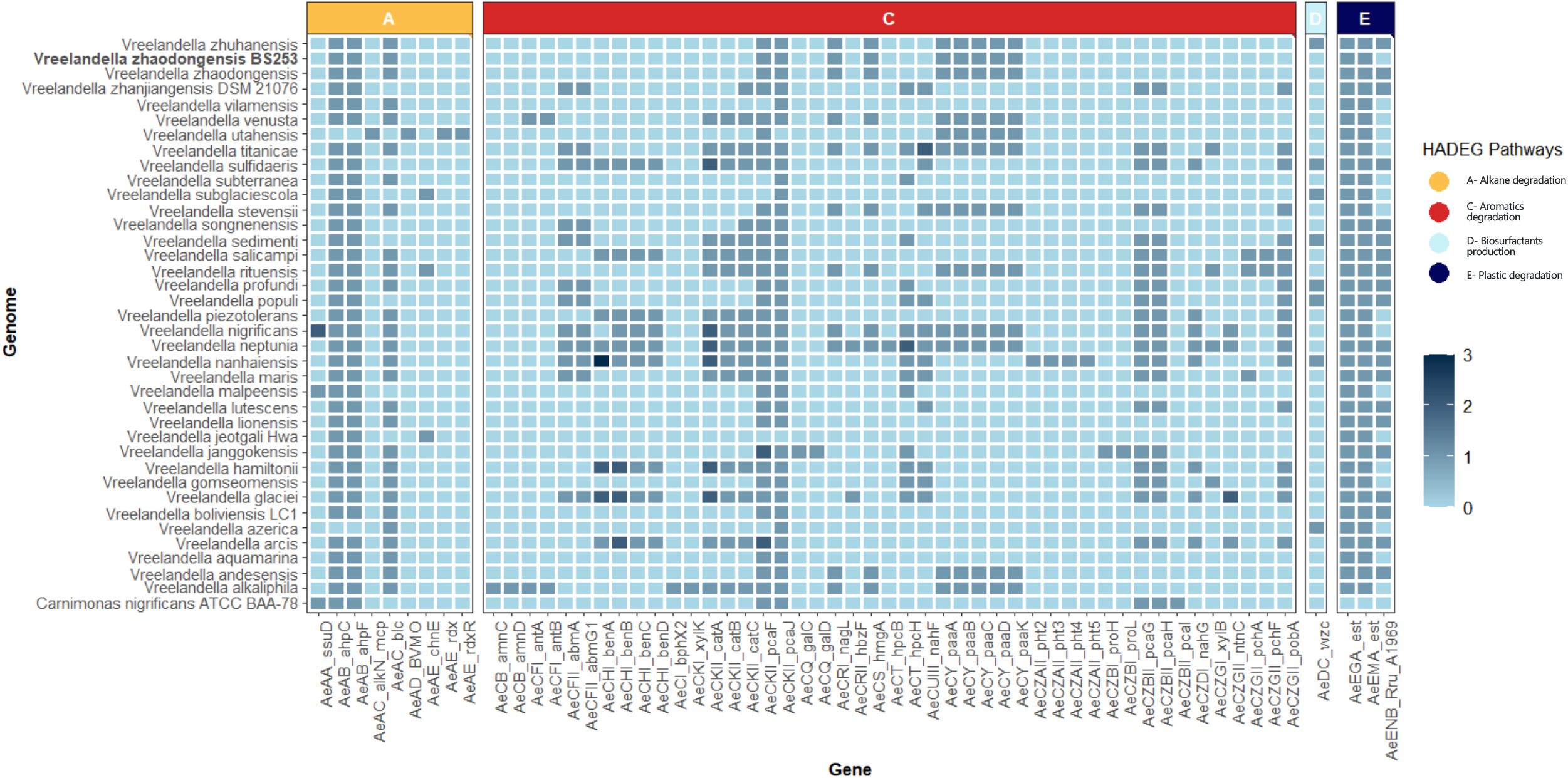
Pathway groups related to alkanes, aromatics, and plastic degradation. Heatmap with the validated genes and the number of hits identified in isolate BS253 and within *Vreelandella*. Gene groups are divided by degradation pathways (A = Alkanes degradation; C = Aromatics degradation; D = Biosurfactants production; E = Plastics degradation). Isolate BS253 and *V. zhaodongensis* (GCF_013415115.1) are highlighted.

The BS253 showed the presence of diverse gene pairs encoding TA systems. We identified a total of 23 pairs from the type II TA system (Additional file: Figure S3 Table S3). The top hits comprised: addiction module antidote protein (*PumPB*) and type II toxin-antitoxin system RelE/ParE family toxin (*PumpA*), type II toxin-antitoxin system VapC family toxin (*VapC*) and plasmid stabilization protein (*VapB*), type II toxin-antitoxin system Phd/YefM family antitoxin (*relB*) and type II toxin-antitoxin system RelE/ParE family toxin (*relE*).

The strain BS253 exhibited the ability to produce potential biosurfactants under the tested conditions. Following cultivation in MSM medium supplemented with 2% glucose, biosurfactant activity was confirmed through emulsification, drop collapse, and oil spreading assays. In the emulsification assay, strain BS253 achieved an emulsion formation rate of 51.7% (Figure 5). As expected, no emulsification was observed in the negative control, while the positive control (0.5% SDS) showed pronounced activity, confirming the reliability of the experimental setup. In contrast, both the drop collapse and oil spreading assays produced negative results.

**Figure 5.**
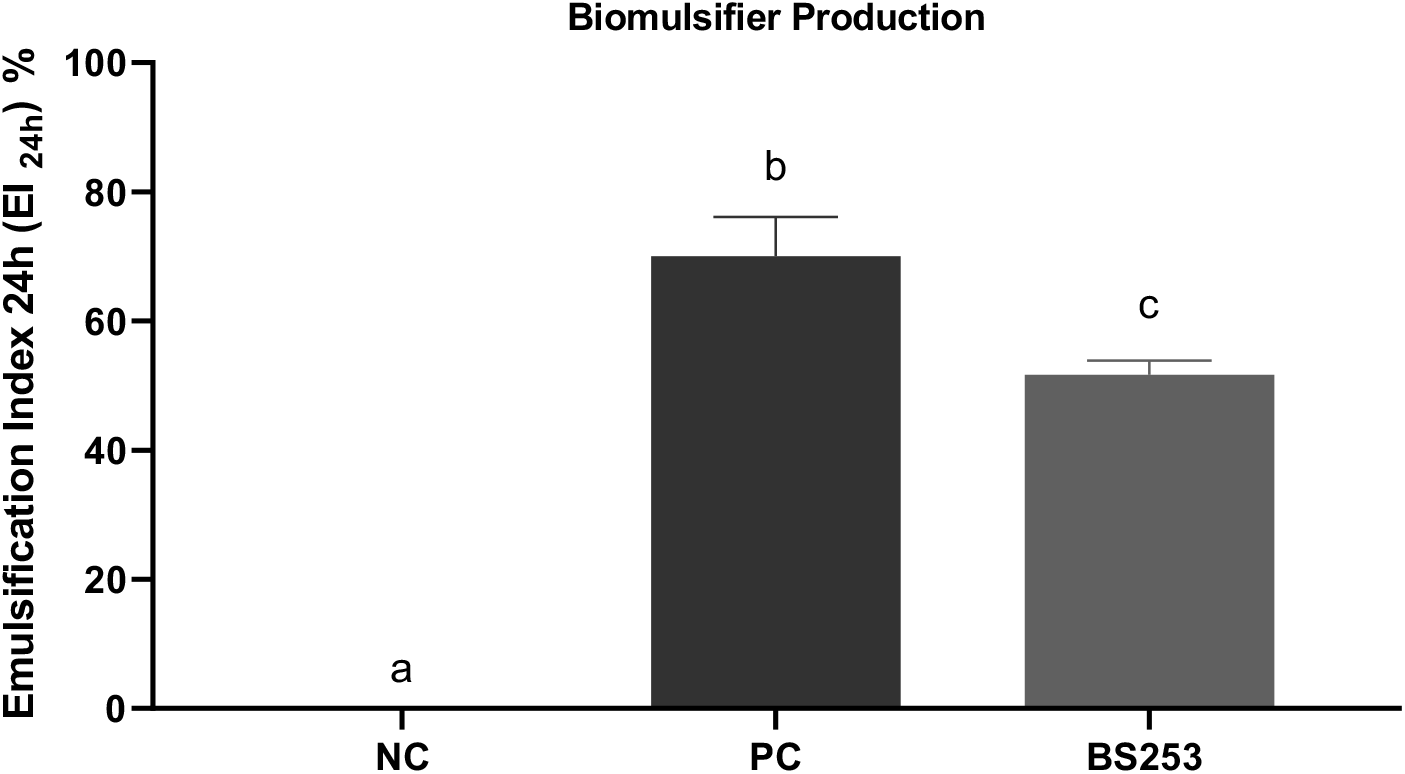
Emulsification index of Vreelandella zhaogonensis strain BS253 cultured in MSM medium supplemented with 2% glucose. The assay was performed using whole-cell cultures after 5 days of incubation. NC: negative control (uninoculated culture medium); PC: positive control (0.5% SDS). Different letters indicate statistically significant differences (p < 0.05) according to one-way ANOVA followed by Tukey’s post hoc test.

## Discussion

Natural environments are home to large assemblages of organisms that evolved to solve diverse pressures and constraints. Sequencing the genome of environmental microorganisms offers a tool to uncover these unique biological properties and pave the way for the development of new biotechnological applications [15]. In this study, we characterized the first complete genome of a new strain of *Vreelandella zhaodongensis* isolated from a hypersaline alkaline lake in Brazil’s Pantanal using hybrid sequence technologies (Illumina and Oxford Nanopore). Along with this, we conducted a complementary analysis to assess genetic content, biosynthetic capacity for metabolic characteristics of this isolate.

The genus *Vreelandella* belongs to the family *Halomonadaceae*. This genus was proposed by de La Haba *et al*. in 2023 from a taxonomic re-evaluation of the phylogenetically heterogeneous *Halomonas* [13]. Using a multi-marker phylogenetic placement method to contextualize BS253 within *Vreelandella* and supported overall genome-related index, BS253 *de novo* assembled genome best matched *V. zhaodongensis*. The phylogenetic tree confirmed these results and placed BS253 with *V. zhaodongensis* and confirmed the phylogeny structure proposed by La Haba *et al*. in 2024, with *V. alkaliphila, V. Venusta*, and *V. andesensis* as the nearest related species. Interesting, although genetically close to *V. zhaodongensis*, the dDDH value between the reference and BS253 strongly supports a subspecies of *V. zhaodongensis* (dDDH = 71,4%), as previously studies indicated that a value of 79-80% dDDH is a reference for delineating a subspecies [49, 61]. Currently, there are only two deposited genomes for this species on the NCBI genome database: one reference, from the study that described *V. zhaodongensis* (ASM1341511v1) [60], and another deposited by the Global Catalogue of Microorganisms (GCM) 10K type strain sequencing project (ASM4266499v1) [62]. Both genomes are assembled in contig level, with a total of 25 scaffolds. Both studies used Illumina MiSeq and HiSeq short-reads platform to sequence the reference genome and implemented an assembled method developed to this technology. Assemblies using these technologies often produce small and fragmented contigs [30, 63]. In contrast, long reads, such as sequences generated by Oxford Nanopore Technology, produce sequences that are thousands of bp in size, which overlap most of the repeats and, consequently, result in larger and complete contigs [27]. Therefore, due to technical limitations, no study had assembled chromosome-level genomes for *V. zhaodongensis*.

To generate high-quality assemblies, new hybrid approaches have been developed and are now considered the gold standard. Recently, an innovative pipeline, called *Trycycler*, takes advantage of the quality of different assemblers and the polishing step using short reads to generate *de novo* assemblies [25]. Basically, this pipeline uses input independent assemblies, all using long reads, derived from different assemblers to generate a consensus contig sequence. After this step, two additional rounds of polishing can be performed: polishing to remove long-read errors or polishing to remove long-read errors + polishing with short-reads. Using this approach, this method guarantees a final sequence without large and small-scale errors. The genome of BS253 was sequenced and assembled by combining high-coverage Nanopore reads, and high-coverage Illumina reads from the same genome using the steps in the *Trycycler* pipeline. We assembled a gap-free, highly accurate, complete genome with a 3,765,793 bp circular chromosome. Therefore, in this study, we present the BS253 assembled genome as the first complete DNA sequence for *V. zhaodongensis*. This will provide the research community with additional information for this distinct halophilic bacterial group.

Members of *Vreelandella* have been found in a wide range of pH and diverse saline habitats. Close phylogenetic relatives of strain BS253 have been recovered from tropical coastal water, salt pools, saline lakes, and saline-alkaline soils. A common physiological trait among *Vreelandella* species is the presence of adaptive mechanisms that confer tolerance to saline and hypersaline conditions. The Pantanal region harbors numerous hypersaline and alkaline lagoons that serve as unique ecological niches, supporting a high diversity of extremophilic microorganisms. These environments are reservoirs of microbial genetic and metabolic diversity, providing organisms with adaptations to extreme salinity, pH, and temperature fluctuations, which can be exploited for biotechnological applications [8]

Consistent with this profile, strain BS253 exhibited a physiological requirement for minimum NaCl concentrations and alkaline pH to support microbial growth both at 24 hours and 48 hours of culture, corroborating the environmental conditions of its original isolation site—a hypersaline, alkaline lake in Brazil’s Pantanal biome. Our findings further indicate that *V. zhaodongensis* strain 253 displays a phenotype adapted to moderate temperature conditions, with an optimal growth temperature starting from 20⍰°C. The values observed in this strain are lower than those found in the other strains analyzed. The temperature profile and its halophilic, alkalitolerant behavior indicate that BS253 is adapted metabolically and physiologically to the extreme conditions of saline-alkaline ecosystems.

When compared to its phylogenetically closest relatives, strain BS253 lacks motility and catalase activity—traits commonly present in other *Vreelandella* species—and exhibits distinct metabolic features, including a potential for anaerobic respiration [13]. These differences further emphasize its unique physiological profile and suggest specialized adaptations to microaerophilic or anoxic niches within saline-alkaline environments, supporting that the BS253 strain may represent a distinct subspecies within *V. zhaodongensis*. Furthermore, analysis using the API miniaturized system revealed a high degree of metabolic flexibility, particularly in acid production, which further contributes to its distinct phenotypic profile in the BS253 genome.

Halophilic and halotolerant bacteria are reported as potential non-model microorganisms adapted to salt conditions and producers of high-value molecules, including biosurfactants and compatible solutes [64]. Biosurfactants are molecules produced by microorganisms from various carbon sources, with applications in bioremediation and petroleum recovery. In this context, microbial biosurfactants can be obtained by relatively simple procedures, such as fermentation processes involving sugars, oils, alkanes, industrial and agricultural waste. They exhibit diverse chemical structures and surfactant properties, serving various natural functions with distinct applications [65]. To evaluate the potential induction of biosurfactant production by BS253, a screening was conducted, and the result obtained for emulsion formation was 51.7%. The results related to emulsion formation indicate that strain BS253 can synthesize bioemulsifying compounds when grown in MSM with glucose as the sole carbon source. However, the absence of activity in the drop collapse and oil spreading assays suggests that the compound produced may have limited ability to reduce surface or interfacial tension. The exclusive detection of biosurfactant production in MSM suggests that strain BS253 is capable of efficiently uptake and metabolizing glucose even under nutritionally limited conditions. In contrast, no growth was observed in the API® AUX Medium.

Despite being a complex medium designed to assess the carbohydrate assimilation profile of microorganisms, API^®^ AUX contains a mixture of carbon and nitrogen sources that are pre-formulated and not specifically optimized for biosurfactant production (bioMérieux). The lack of growth may be attributed to the absence of essential micronutrients, unfavorable pH, or an imbalance in nutrient ratios that do not support the metabolic requirements of BS253.

Moreover, the presence of multiple carbon sources in the API^®^ AUX Medium could lead to catabolite repression, a regulatory mechanism that inhibits the uptake and metabolism of certain substrates—such as glucose—when others are available in excess [66]. In contrast, MSM is a minimal, well-defined medium with a single carbon source, allowing for more targeted metabolic activation [67, 68]. The ability of BS253 to grow and produce biosurfactants in MSM reinforces its metabolic selectivity and suggests that streamlined nutrient conditions may be more effective in triggering secondary metabolism and biosynthetic pathways involved in biosurfactant synthesis. These findings reinforce the strain potential for bioproduct generation and support its applicability in biotechnological processes focused on biosurfactant production, particularly those based on low-cost, defined carbon sources.

*Halomonas* spp. produce a variety of biotechnologically useful molecules, such as lipids, enzymes, and polysaccharides, that can be employed in several industrial processes [69]. Among them, increased research interest highlighted the capacity to produce biosurfactants and bioemulsifiers by *Halomonas*. Studies have demonstrated that among oil-degrading bacteria and biosurfactant producers, iturin was among the most abundant pathways in a microbial consortium [70]. Iturin A, produced by certain *Bacillus* strains, exhibits strong surface-active properties, making it an effective surfactant and emulsifier. This result supports the view that halophilic and halotolerant bacteria constitute efficient platforms for the biotechnological production of high-value molecules, including biosurfactants.

Strategies that enhance substrate bioavailability, such as biosurfactant production, may increase the biodegradability of aromatic compounds. Numerous microorganisms that break down aromatic compounds have unique physiological mechanisms that enhance the availability of compounds with low solubility in water [70–72].The production of biosurfactants can improve the bioavailability of these substances by either raising their apparent solubility in the aqueous phase or by increasing the contact surface area through emulsification. In our study, we identified in the BS253 the presence of routes for key pathway groups related to Alkanes degradation, Aromatics degradation, and Plastic degradation. In parallel, with an emulsification test, an emulsion index of 51.7% supported the biosurfactant production by BS253.

The biotechnological potential of molecules produced by organism like 253, place them as optimal sustainable candidates over conventional physical and chemical methods used in carbon capture approaches. Recent assessments by international agencies such as the IPCC and IEA emphasize the urgency of developing scalable and efficient CCUS technologies to meet global climate targets [73, 74]. In this context, previous studies have underscored that *V. stevensii* can grow in CO_2_-rich streams and demonstrated substantial remove CO_2_ from the medium [75, 76]. This sustain the capacity of this clade be used in biotechnological strategies to carbon capture and fixation in high value molecules.

The identification of strains like *V. zhaodongensis* BS253, capable of withstanding hypersaline and alkaline stress while producing biosurfactants under defined culture conditions, offers new opportunities for expanding the repertoire of microbial platforms suitable for CCUS. As demonstrated in this study, the integration of genomic, metabolic, and physiological analyses provides a foundation for selecting and engineering extremophilic strains with enhanced biosurfactant production capacities. These insights contribute to the development of biobased CCUS systems aligned with circular economy principles and sustainable industrial practices.

## Conclusions

In this study, we present the first complete genome sequence of *V. zhaodongensis* strain BS253, isolated from a hypersaline alkaline environment in the Brazilian Pantanal. Through comprehensive genomic, phylogenomic, and phenotypic analyses, we establish BS253 as a distinct subspecies within *V. zhaodongensis*, with notable adaptations to extreme saline-alkaline conditions. The strain exhibits a unique combination of physiological traits and genomic features, as well as genes associated with hydrocarbon and plastic degradation pathways. The confirmed biosurfactant activity, along with the strain’s ability to thrive in defined, minimal media, underscores its potential as a microbial platform for sustainable biotechnological applications.

This work expands the genomic resources for the *Vreelandella* genus and highlights the untapped potential of extremophiles from underexplored ecosystems. The isolation of microbial organisms from extreme environments is pivotal for the identification of new strains with potential for ecological and biotechnological applications to use in CCUS strategies [4, 5]. Therefore, our metabolic and genomic results suggest that the isolate BS253 offers promising capabilities for bioeconomy development.

## List of abbreviations

## Declarations

Not applicable

### Ethical Compliance

All procedures performed in studies involving human participants were in accordance with the ethical standards of the institutional and/or national research committee and with the 1964 Helsinki Declaration and its later amendments or comparable ethical standards.

### Consent for publication

Not applicable

### Availability of data and material

The dataset generated and analyzed during the current study are available in the BioProject ID PRJNA1275882 (https://www.ncbi.nlm.nih.gov/bioproject/PRJNA1275882).

### Authors’ contributions

## Acknowledgment

We thank the staff of the Life Sciences Core Facility (LaCTAD) from the State University of Campinas (UNICAMP) for the Nanopore Sequencing services. The authors thank the Institute of Petroleum and Natural Resources (IPR) of the Pontifical Catholic University of Rio Grande do Sul for the infrastructure. We also acknowledge the use of *parallel* software [77].

