## Supplemental file 1 for "Complete genome sequence and metabolic features of *Vreelandella zhaodongensis* BS253: A new isolate from hypersaline lakes from Brazilian Pantanal"

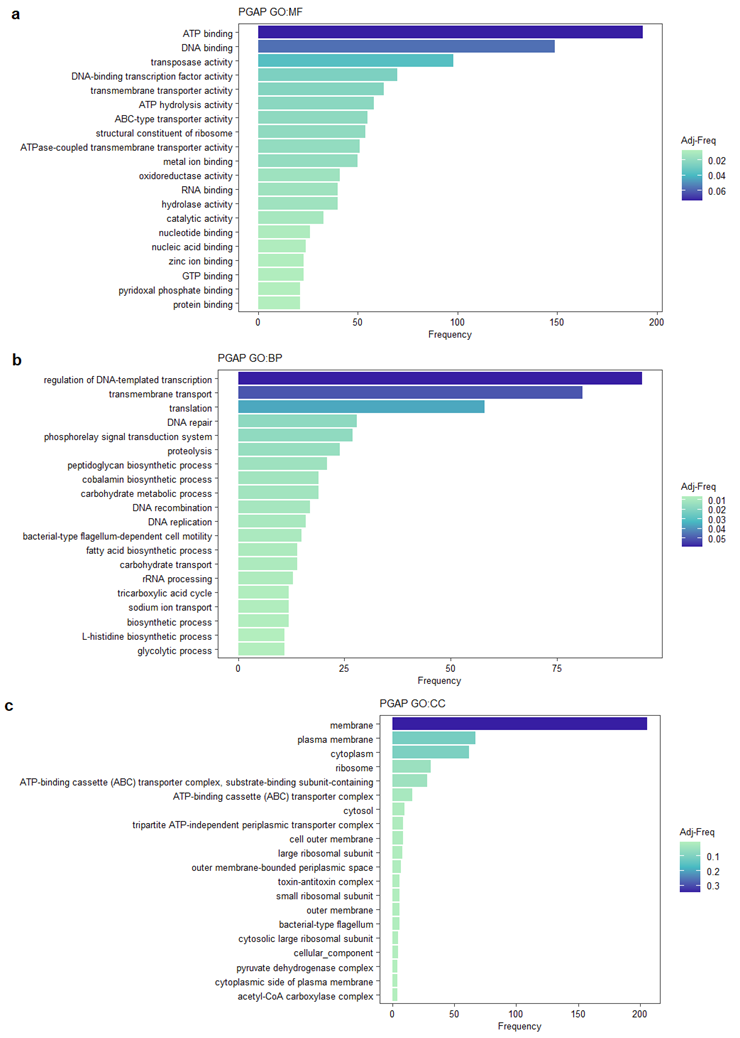

**Figure S1.** **Functional annotation of the complete *de novo* assembled genome of *V. zhaodongensis* BS253. (A-C)** Top 20 most represented GO functional annotations annotated using PGAP. Panels represent GO categories for Molecular Function (MF, Panel A), Biological Process (BP, Panel B), and Cellular Component. Each category is described on the y-axis, while the number of genes in each category is displayed in horizontal bars along the x-axis. The bars are colored according to the percentage of each category (Adj-Frequency, Adj-freq.

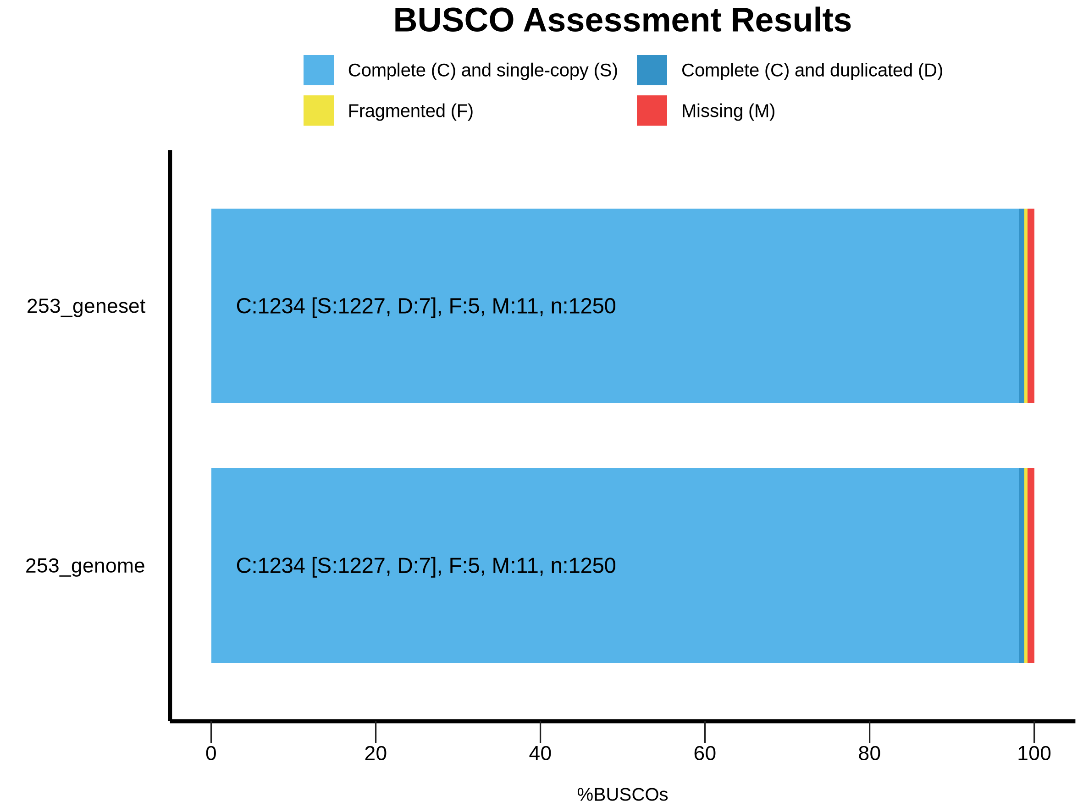

**Figure S2. BUSCO evaluation results for the de novo assembled genomic sequence and the annotated gene set.** BUSCO tool v5.8.2 using the specific reference halomonas_odb12 (Creation date: 2024-11-14, number of genomes: 102, number of BUSCOs: 1250). Gene set 253 (C:98.7% [S:98.2%, D:0.6%], F:0.4%, M:0.9%, n:1250) and genome 253 (C:98.7% [S:98.2%, D:0.6%], F:0.4%, M:0.9%, n:1250).

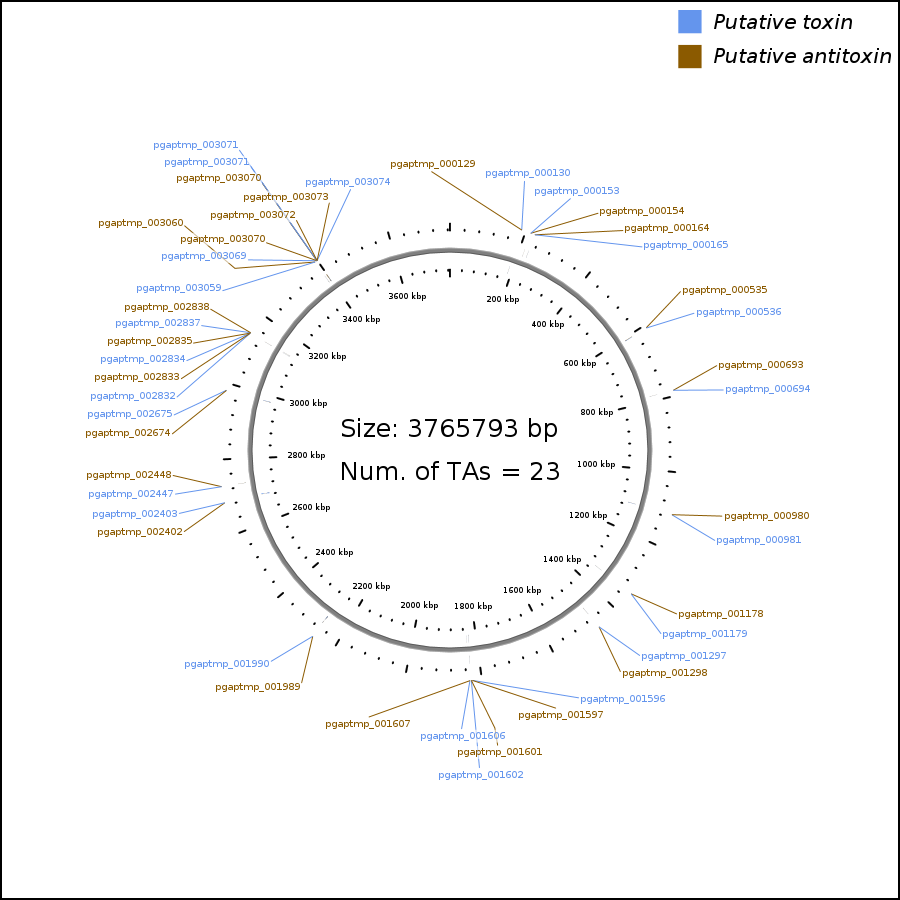

**Figure S3. Toxin-antitoxin pairs in the genome of isolate 253.** The annotated genome showed 23 putative TA pairs annotated in the TADB database. Blue, putative toxin genes; Brown, putative antitoxin genes. Generated using *TAfinder* 2.0 (https://bioinfo-mml.sjtu.edu.cn/TADB3/TAfinder.php).
